# Genomic landscape, polymorphism and possible LINE-associated delivery of G-Quadruplex motifs in the bovine genes

**DOI:** 10.1101/2021.12.13.472480

**Authors:** Georgios C. Stefos, Georgios Theodorou, Ioannis Politis

## Abstract

G-Quadruplex structures are non-B DNA structures that occur in regions carrying short runs of guanines. They are implicated in several biological processes including transcription, translation, replication and telomere maintenance as well as in several pathological conditions like cancer and thus they have gained the attention of the scientific community. The rise of the –omics era significantly affected the G-quadruplex research and the genome-wide characterization of G-Quadruplexes has been rendered a necessary first step towards applying genomics approaches for their study. While in human and several model organisms there is a considerable number of works studying genome-wide the DNA motifs with potential to form G-quadruplexes (G4-motifs), there is a total absence of any similar studies regarding livestock animals. The objectives of the present study were to provide a detailed characterization of the bovine genic G4-motifs’ distribution and properties and to suggest a possible mechanism for the delivery of G4 motifs in the genes. Our data indicate that the distribution of G4s within bovine genes and the annotation of said genes to Gene Ontology terms are similar to what is already shown for other organisms. By investigating their structural characteristics and polymorphism, it is obvious that the overall stability of the putative quadruplex structures is in line with the current notion in the G4 field. Similarly to human, the bovine G4s are overrepresented in specific LINE repeat elements, the L1_BTs in the case of cattle. We suggest these elements as vehicles for delivery of G4 motifs in the introns of the bovine genes. Lastly, it seems that a basis exists for connecting traits of agricultural importance to the genetic variation of G4 motifs, thus, cattle could become an interesting new model organism for G4-related genetic studies.

## 1. Introduction

There are specific nucleotide sequences that under certain conditions do not adopt the right-handed Watson-Crick B-form but instead form non-B DNA secondary structures like G-quadruplex, Z-DNA and cruciform (Wells et al., 2007(1)). The G-quadruplex structure occurs when short runs of guanines (Gs) form G-quartets held by multiple Hoogsteen hydrogen bonds and are stacked one over the other stabilized by cations (Spiegel at al., 2020(2)). There is a variety in G-quadruplexes’ structure which arises from factors like whether Gs come from one or more DNA strands, the orientation of the strand, the number of G-runs, whether the structure is adopted by a DNA or an RNA molecule and others (Burge et al., 2006; Mukundan et al., 2013; Malgowska et al., 2016(3–5)).

The most well-recognized nucleotide sequence pattern for G-quadruplexes is (G_≥3_N_1-7_)_≥3_G_≥3_, meaning at least four G-runs interrupted by loops that have length from one to seven nucleotides. One of the most used algorithms for scanning DNA sequences for the above pattern is Quadparser (Huppert and Balasubramanian, 2005(6)). However, more sophisticated algorithms have been also developed which are more flexible in terms of number and size of G-runs and loops, allow mismatches, identify patterns involving more than one strands or even algorithms that are trained by experimentally validated data (Lombardi and Londono-Vallejo, 2020(7)). The occurrence of the motifs identified by the aforementioned algorithms (called G4 motifs hereafter) does not necessarily mean the formation of G-quadruplex structures. For the genome-wide experimental validation of the existence of G-quadruplex structures several methods have been developed that can be applied in both DNA (Lam et al., 2012; Rodriguez et al., 2012; Hänsel-Hertsch et al., 2016; Yoshida et al., 2018; Marsico et al., 2019; Zeng et al., 2019(8–13)) and RNA molecules (Kwok et al., 2016; Yang et al., 2018(14,15)).

G-quadruplexes are implicated in several biological processes like replication, telomere maintenance and protein expression among others (Prioleau et al., 2017; Bryan, 2020; Varshney et al., 2020(16–18)). Their crucial role in expression is underlined by their high occurrence in genes and has been shown by both experimental and bioinformatics approaches. Thus, G-quadruplexes at the promoter regions can regulate transcription by affecting the recruitment of the transcriptional machinery (Rigo et al., 2016(19)). When located on the genic regions *per se*, G-quadruplexes can facilitate or repress the transcription by RNA polymerase, depending on which strand they are located on. When they are on the template strand they have a negative effect on transcription by acting as obstacles for RNA polymerase, while when on the coding strand they can have either a positive effect by maintaining DNA in an open state or a negative effect by favoring the formation of DNA:RNA hybrids (Varshney et al., 2020(18)). In addition to transcription, G-quadruplexes that are formed on the RNA molecules can regulate translation. Thus, RNA G-quadruplexes located on 5’UTRs affect translation initiation by facilitating recruitment to ribosomes while those located on the CDSs and 3’UTRs play rather an inhibitory role in translation. Several other roles have been attributed to genic G-quadruplexes like RNA localization, polyadenylation, splicing and epigenetic regulation, each of them with an impact on protein expression (Varshney et al., 2020; Song et al., 2016(18,20)).

During the last decade there is a highly increased interest for G4-quadruplexes. This interest is derived from the association of these structures with several fundamental biological processes as well as with several diseases including cancer (Nakanishi et al., 2020; Dumas et al., 2020; Varshney et al., 2020(18,21,22)). The rise of the -omics approaches, that also took place during the last years, is reflected in the G-quadruplex research. Plenty of proteomics and genomics studies have been conducted towards a comprehensive understanding of the different roles that these structures play. In parallel, studies following data-driven approaches are conducted while all the above are supported by the development of new bioinformatics tools.

Although the importance of rodents as model organisms in biomedical and translational research is crucial and unquestionable, the use of livestock animals as alternative/parallel models has major advantages. Specifically, livestock animals present several similarities in anatomy, physiology, metabolism as well as in the organs size with humans while in many cases the pathogenesis of diseases is closer to that of humans (Roth et al., 2015; Smith and Govoni 2022(23)(24)). Furthermore, livestock animals constitute an invaluable resource for genetics research to gain insight into genotype-phenotype relationships, given their rapid evolution by selection since their domestication (Andersson 2013(25)). New sequencing technologies have been used extensively during the last few years in cattle genetic/genomic research. A considerable and rapidly increasing number of bulls have been subjected to whole genome sequencing while phenotyping/genotyping is being done in an industry-scale level with millions of animals being genotyped with SNP chips with comprehensive phenotype recording in pedigrees. The rapid identification of mutations linked to sporadic syndromes that also occur in humans, also renders cattle as attractive models (Hayess et al., 2019(26)). In parallel, efforts like the 1000 bulls genome project or the Functional Annotation of Farm Animal Genomes (FAANG) for comprehensive mapping of functional/regulatory elements genomewide, support cutting-edge strategies in the postgenomic era (Hayess et al., 2019; Tixier-Boichard et al., 2021(26,27)). Indeed, there are several examples that the above-mentioned advances in cattle genetics have allowed this species to successfully model diseases and traits (Bouwman et al., 2018; Bourneuf et al., 2017(28,29)). Despite the ongoing advances in genomics and genetics of livestock animals and especially cattle, there is a gap in the G4-related studies and an absolute absence of genomewide studies for this species (Stefos et al 2021(30)). Efforts to study G4s in cattle could have double benefit in both biomedical and farm animal research.

Here we studied the distribution of G4-quadruplex motifs that are associated with genes and promoters in the genome of Bos taurus. We characterized in detail G4 motifs in promoters, introns, coding and untranslated regions and we delineated the polymorphisms associated with these regions. Finally, we considered the co-localization of G4-motifs and repeat elements and we suggested the LINE family L1_BT as a vehicle for G4-motif distribution within introns. To the best of our knowledge this is the first genome-wide study regarding the G4-motifs of a livestock species.

## 2. Material and methods

### 2.1. Genomic data acquisition

Coordinates of bovine G4 quadruplex motifs (assembly: bosTau3) were downloaded from the Non-B DB database (Cer et al., 2013(31)). The coordinates of the bovine repeat elements (assembly: bosTau8) (LINE, SINE and LTR) and the SNPs (assembly: bosTau8) from dbSNP build 148 (SNPs148) were downloaded from the UCSC Table browser. SNPs from chr17 were not accessible at the time of data analysis and thus they have not been included in the analyses. The BGVD SNPs (assembly: bosTau5) were downloaded from the Bovine Genome Variation Database (Chen et al., 2020(32)) while the SNPs of the BovineHD BeadChip (assembly: bosTau4) were obtained from Illumina’s site. The QTL/association data (assembly: bosTau6) were obtained from the Cattle QTLdb (www.animalgenome.org) (Hu et al., 2018(33)). QTLs and associations differ in a) the methods they are identified with and b) their chromosomal resolution, as it is described in the site of the database. Where necessary, all the aforementioned data were converted to Bos_taurus_UMD_3.1.1/bosTau8 assembly coordinates using the UCSC Genome Browser LiftOver tool.

### 2.2. Feature annotation

The annotation of the G4 motifs on the bovine genic elements was done using the annotation of UCSC RefSeq and the findOverlaps function of GenomicRanges in R (Lawrence et al., 2013(34)). G4 motifs overlapping with more than one genic features (of the same or different genes) were assigned to all of them. As template strand is defined the one that RNA polymerase is bound on and as coding strand the one that has the same sequence with the transcribed RNA. In the case of promoter regions the terms “coding” and “template” may have not any practical meaning, even though it is used for enabling the examination of the strand-related bias.

### 2.3. G4 characterization

The characteristics of G4 motifs like number and lengths of G-runs and loops per motif were calculated using a custom R script (supplementary file 1). The DNA sequences of G4-motifs were retrieved using the getSeq function of Biostrings and the BSGenome object for Bos_taurus_UMD_3.1.1/bosTau8. When the orientation of the motifs was taken into account, as first G-run (or loop), the most upstream one according to the direction of transcription was defined. Only the G4 motifs with a number of nucleotides smaller or equal to 110 were taken into account.

### 2.4. Ontology terms overrepresentation analysis

Ontology overrepresentation analyses, for both Gene Ontology and Panther Protein Class, were done using the online tool Panther (Mi et al., 2010(35)). As background list, all bovine genes were used. P-values that are shown have been filtered for FDR < 0.05. “Counts” indicates the number of genes that carry G4s on their feature of question and are annotated to the relative ontology term. “Term size” indicates the number of bovine genes that are annotated to each ontology term regardless if they carry G4 motifs or not.

### 2.5 G4 motifs on human orthologues

The human orthologues of the bovine genes that carry G4s were obtained using bioDBnet (Mudunuri et al., 2009(36)). The fasta files with the DNA sequences of the genic features of these genes for the hg38 were downloaded from BioMart. The fasta files were analyzed with pqsfinder (Hon et al., 2017(37)) for the presence of G4s motifs with the same pattern as those downloaded from the Non-B DB database.

### 2.6 Analysis of disruptive Gs

For the identification of SNPs on disruptive Gs, the procedure below was followed. First the coordinates of all Gs within G-runs for each genic feature that meet the criteria to be disruptive according to Nakken et al., (2009(38)) were found. This was done using the matrix with the number/lengths of G-runs and loops of all G4s that is produced with the custom R script provided in the supplementary material. Briefly, as disruptive were assigned the following Gs: all Gs of all 3 nt-long G-runs, the 2^nd^ and 3^rd^ Gs of all 4 nt-long G-runs and the middle G of all 5 nt-long G-runs. Next, overlaps of these disruptive Gs with the SNPs of the tested databases/arrays were identified using the function mergeByOverlaps of the R package GenomicRanges.

### 2.7. L1_BT repeats alignment

The alignment of the 1000 random intronic L1_BT repeats with a full-length element (GenBank: DQ000238.1) was done using the pairwiseAlignment function from the Biostrings R package using type of alignment: local.

### 2.8. Statistical analyses and plots

The statistical analyses used in the present study were conducted in R and are thoroughly described in the figure legend of each plot. Plots were made either in R or Excel.

## 3. Results and Discussion

### 3.1. Distribution of G4s on bovine genic features

The predicted G4 motifs of the bovine genome were downloaded from the Non-B DB database (Cer et al., 2013(31)). The G4s of this database were identified by an algorithm that scans for motifs of the following pattern: (G_≥3_N_1-7_)_v≥3_G_≥3_. This formula is considered rather conservative since other algorithms, as well as experimental methods, can predict the existence of more motifs (Hänsel-Hertsch et al., 2017(39)). However, the above formula when validated via the performance of Quadparser, the informatics tool that first used it, although it was characterized as not sensitive, it was considered specific (Lombardi and Londono-Vallejo, 2020(7)).

The density of the G4 motifs that met the above criteria in the bovine genome is 132 G4s/Mb. This density is close to that of human and horse and in the same range with other mammals (mouse and dog) and it is considerably different to that of arabidopsis (Fig. 1A). The analyses that will be reported hereafter will concern the motifs that fall within genic features, namely 5’-UTRs, CDSs, introns and 3’-UTRs as well as within the promoter regions that exist 500 bp upstream the transcription start site, named here as prom500. The sum of these gene-associated G4 motifs consists about 30 % of the database’s motifs. The full list of all gene-associated G4 motifs along with information regarding their loci on the bovine genome, the genic element they are located on, the transcripts that they are associated to and the DNA strand relatively to transcription that they are located on, is provided in the Supplementary Table 1. In Fig. 1B the G4 density of each bovine genic feature is shown. The promoters and the untranslated regions are the most enriched among the genic features and are by far more enriched than the genome’s average density. Elevated density in UTRs is also seen in predicted G4s in humans while this does not seem to be the rule for plants (Kopec et al., 2019(40)). Higher density in UTRs compared to CDSs has been also reported for experimentally observed RNA-G4s in humans (Kwok et al., 2016(14)). When considering genome-wide experimentally observed G4s on DNA, promoters and 5-UTRs are clearly enriched compared to the genome in both humans and mice, while 3-UTRs are slightly enriched in human but not in mouse (Marsico et al., 2019.pdf(41)).

**Figure 1:**
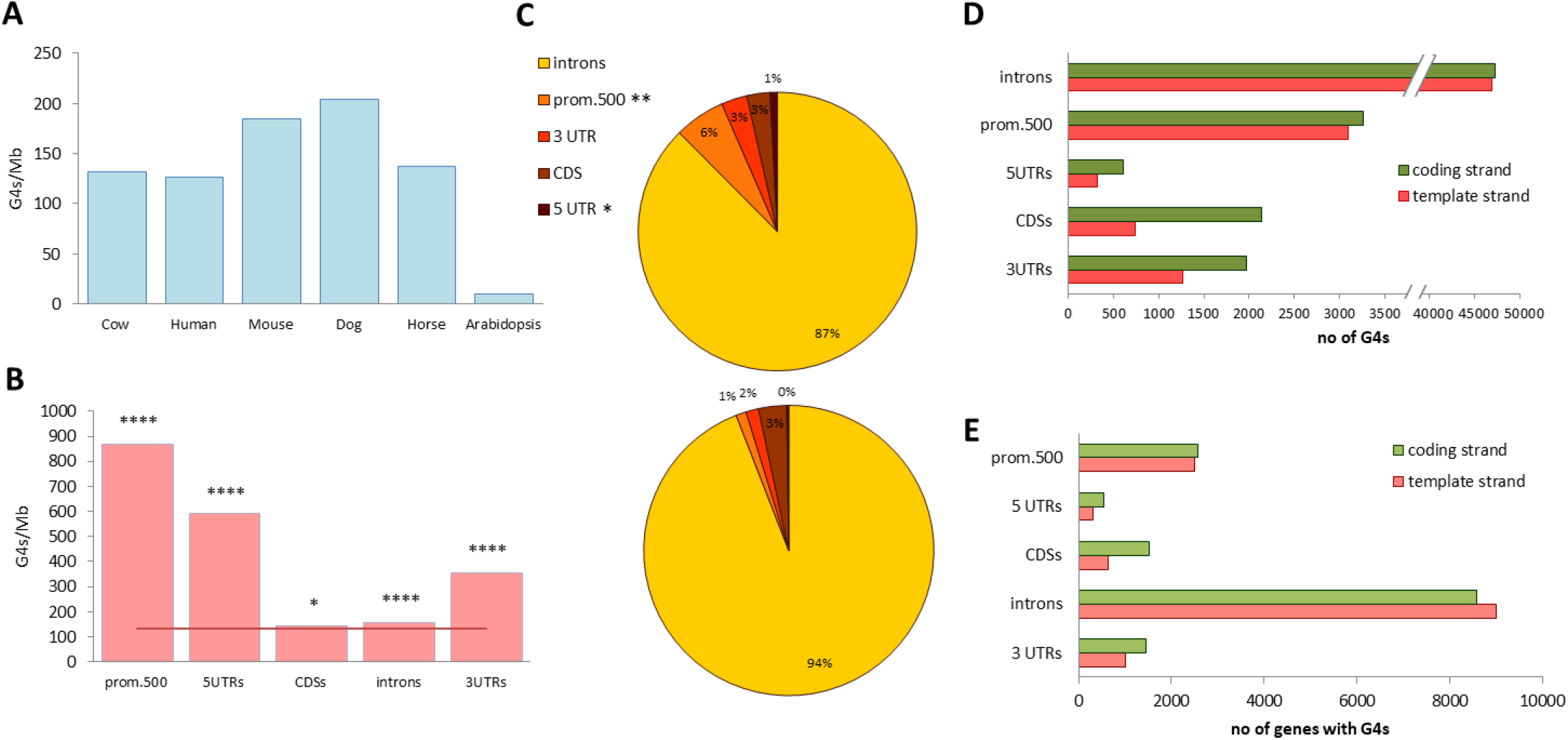
Occurrence of G4 motifs on cattle genes. **A**. Density of G4 motifs in several species. Densities were calculated based on the number of G4s deposited in the non-B database for each species and the total ungapped length of the genome assemblies that were used in the database. **B**. Densities of G4 motifs within the bovine genic features. The densities were calculated by dividing the number of G4s annotated to each feature by the feature’s total length. The horizontal red line indicates the whole genome’s density. The asterisks indicate statistical difference between the features’ and whole genome’s densities. The difference was tested by comparing each feature’s density to the density of areas consisting of 2000 random genomic intervals of length equal to the mean length of the feature. **C**. Distribution of G4 motifs on bovine genic features. Upper pie chart: Actual proportions of G4 motifs annotated on bovine genic features. The asterisks indicate statistical difference between the actual proportions and the ones that resulted by randomization tests. In the randomization tests the expected proportion was calculated based on the number of G4s in random 500 nt long fragments of total length equal to the 1% of each genic feature’s actual total length; Lower pie chart: Expected proportions of G4s on genic features based on each feature’s length. Proportions are rounded for the sake of visualization. **D**. Number of G4 motifs located on coding and template strands of bovine genic features. **E**. Number of genes that carry G4 motifs on their genic features. One gene may carry more than one motif on the same feature or can have G4 motifs in more than one of its features. For the statistical tests, 1000 randomization tests were conducted for each genic feature. For the sampling of the random intervals the ambiguous bases (Ns) were masked. As p we defined the ratio of the randomization tests with different (higher or lower) than the expected value *, p < 0.05; **p < 0.005; ****, p < 0.0001.

The actual proportions of G4s per genic feature can be seen in Fig. 1C (upper pie chart). The largest part of G4 motifs is annotated to introns, which is expected given that the latter has by far the biggest size among the features. When comparing the actual and the expected G4 proportions, it is clear that G4 motifs are enriched in promoters and 5’-UTRs while they tend to be depleted in the introns (p=0.067). The lower panel of Fig. 1C depicts the expected proportions according to the features’ total lengths.

Fig. 1D shows the G4 distribution in the coding and template strand for all genic elements. There is an obvious difference between the two strands only for the exonic features and this difference cannot be credited to G4 occupancy bias between the two DNA strands of the bovine genome (175,303 and 176,139 motifs for the two strands). The attribution of certain functional roles on G4s when present on mRNA molecules could explain the exon-specific elevated G4 occurrence on the coding strand (Varshney et al., 2020(18)).

Concerning the G4s on 5-UTRs, the number of those found on the coding strand is almost double of those on the template strand. Such a strand asymmetry is also reported for the human genome and is suggested to be linked to regulation of translation initiation (Huppert et al., 2008(42)). The same strand asymmetry has also been reported for plants (Busra Cagirici et al., 2020(43)). A bias in strand occupancy is also clear for the 3-UTRs. In humans the strand bias in 3-UTRs is suggested to be linked to effective cleavage at polyadenylation sites (Huppert et al., 2008(42)). Among all genic features, CDSs are those with the highest difference between the two strands (almost 3 times difference). Given that the numbers of expected and observed G4s on CDSs are similar (Fig. 1C), the strand bias in this feature could result from both an overrepresentation of G4s in the coding strand and an underrepresentation in the template strand. Introns do not show any noticeable difference between the two strands, however the coding strand has slightly more motifs. Among the suggested functions of the intronic G4s are transcription elongation and RNA splicing (Varshney et al., 2020(18)). While the former seems to be associated with G4s of both DNA strands, the latter seems to depend on G4s of the coding strand that bind splicing proteins. According to a study not peer-reviewed yet, such G4s are overrepresented near the edges of the introns (Georgakopoulos-Soares et al., 2019 PREPRINT(44)). Given the considerable total length of the introns, these G4s render a small fraction of the total intronic G4s. The small difference of the coding strand G4s over the template strand (which in absolute numbers is significant) could reflect the enrichment of these splicing-associated G4s. The absence of any strand asymmetry in the bovine promoters that we observe (Fig. 1D) is in line with the results of a study by Du et al. (2008(45)). The authors show that in the region 500 bp upstream the TSSs in five animal species there is no difference in G4 occupancy between the two DNA strands. The G4s in the promoter regions have been suggested to interplay with regulation of transcription by enabling transcription factor binding as well as chromatin remodeling (Kim, 2019(46)). Since the transcription factor binding motifs do not seem to have any preferred orientation (Lis and Walther, 2016(47)), the aforementioned proposed roles of the promoter G4s could justify the observed absence of strand bias.

The difference in G4s number between coding and template strands is reflected also in the number of genes carrying G4s (Fig. 1E). There is the exception of introns for which, although there are more G4 motifs in the coding strand, the genes with G4s in the coding strand are slightly less than those with G4s on the template strand.

Fig. 2 shows a selection of interesting Gene Ontology (GO) terms that are enriched in genes with DNA and RNA G4 motifs and are common between cattle and several other eukaryotic organisms, including humans (Huppert et al., 2007; Huppert et al., 2008(42,48); Garg et al., 2016(49); Kwok et al., 2016(14)). These GO terms refer to processes that are already associated with G4s, like transcription and localization, as well as to others that their association with G4s is less studied, like development. Moreover, it should be noted that for both cattle and humans the “*G protein-coupled receptor signaling pathway”* and its relative terms, as well as *ribosome*- and *olfaction*-related terms are underrepresented in genes with G4s in promoters or UTRs (Huppert et al., 2007; Huppert et al., 2008(42,48)). Supplementary Fig. 1 shows that half of the human orthologues to the bovine genes that carry G4s on their genes, also carry G4 motifs. This ratio is lower for the genes with G4s on their promoter regions. This considerable overlap between bovine and human G4-carrying genes could partially explain the commonly overrepresented GO terms. Taken together, the above show that G4 motifs in cattle do not exhibit any unexpected distribution among genes and suggest a compliance of their functional roles with those of other organisms.

**Fig. 2:**
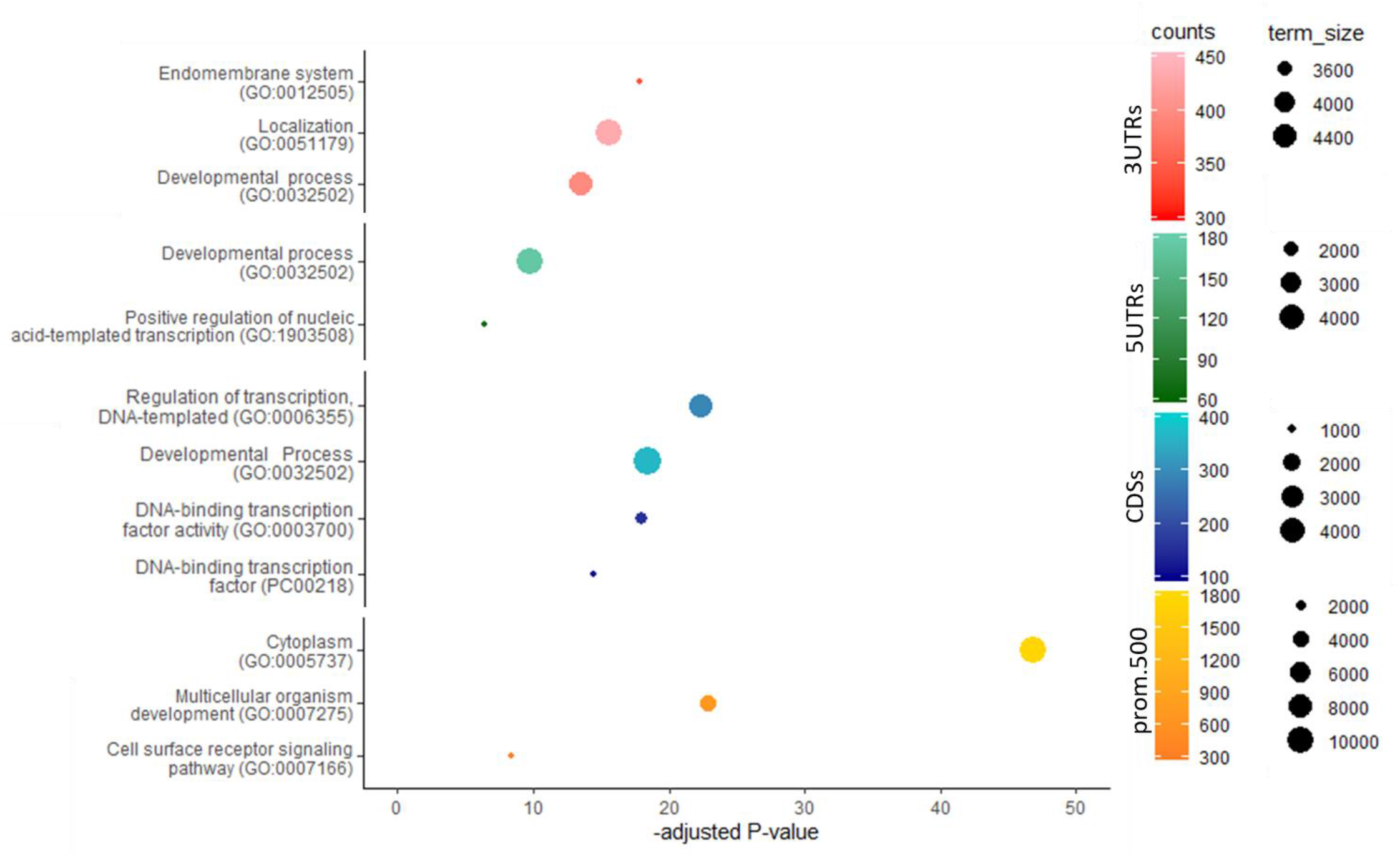
Common ontology terms between cattle and other species enriched in genes that carry G4 motifs on their genic features.

### 3.2. Characterization of G4 motifs on features

The structure of G4 motifs in terms of number and length of G-runs and loops affect their properties including their thermodynamic stability (Lane et al., 2008(50)). Beyond this, functional studies have linked structural properties of G4s to their biological properties (Wieland et al., 2007(51)). Efforts for understanding the role of distribution of the G4 structural patterns in other species have shown that there is bias in the occurrence and distribution for some of them (Huppert and Balasubramanian, 2005; Todd et al., 2005; Garg et al., 2016(6,49,52)). In this context, we decided to analyze in depth the distribution of distinct structural patterns of the bovine G4s motifs on the genic features.

Fig. 3A shows the average number of G-runs per G4 motif for each DNA strand of the studied genic features. While the average number in the genome is around 4.5, the G4 motifs on the promoter regions and on the coding strand of the 5-UTRs have significantly more G-runs. On the other hand, G4 motifs located on the coding strand of CDSs and both strands of introns have less. The density of human G-runs with potential to form G-quadruplex structures as well as plant G4 motifs with diverse number of G-runs are also differentially distributed within genic features and DNA strands, suggesting the existence of distinct functional roles for each G4 type (Eddy and Maizels, 2008; Garg et al., 2016; Mullen et al., 2010(49,53,54)).

**Figure 3:**
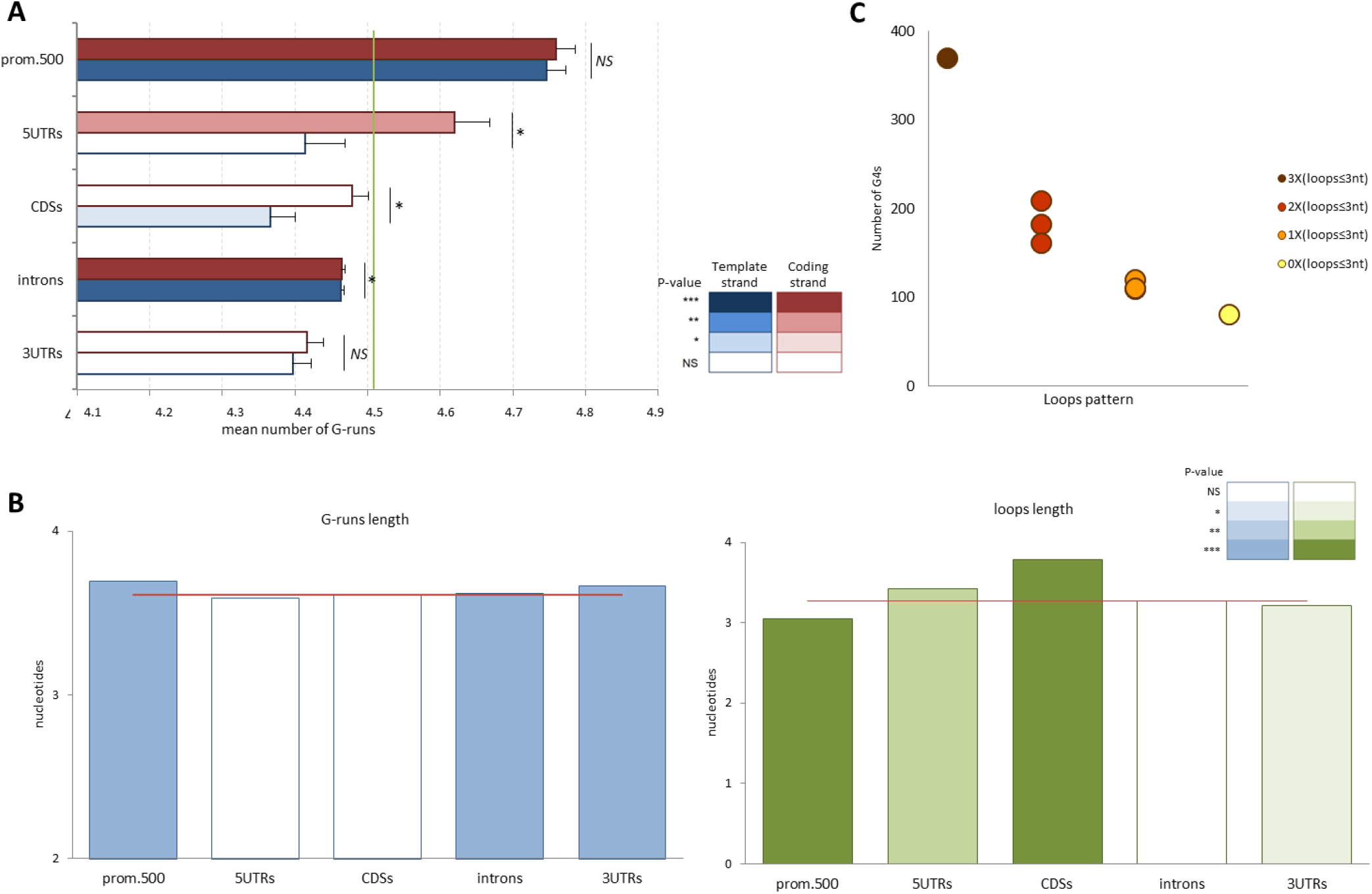
Structural characteristics of bovine genic G4 motifs. **A**. Average number and standard errors of G-runs per G4 motif for all genic features. The color of the bars indicates the significance of the difference to whole genome’s G4s (indicated by a vertical green line). Significance of difference between the two strands within each feature is shown at the right of each bar-couple. Statistical significances were tested by Mann-Whitney test. **B**. Average length of G-runs (left panel) and loops (right panel) of G4 motifs for each genic feature. The horizontal red lines indicate the average length of G-runs and loops in whole genome. For both genic features and whole genome, only the G4s with a total length less than 110 nt and having exactly four G-runs were taken into consideration. The color of the bars indicates the statistical significance between the average length of each feature and that of the whole genome. For calculating this significance the average length of each feature was compared to an equal sized list of G4s picked up randomly from the whole genome. This comparison was conducted 1000 times for each feature. P-value was defined as the ratio of the randomization tests with average size different (higher or lower) from the feature’s average size. **C**. Number of G4 motifs located on both DNA strands on promoters and carry 3, 2, 1, or no loops of length smaller or equal to 3 nucleotides. The G4s with four G-runs and length of G-runs smaller or equal to 4 nucleotides were only taken into consideration. NS, not significant; *, P < 0.05; **, P < 0.005; ***, P < 0.001.

We next examined the length of G-runs and loops given that both affect the stability of the quadruplex structure. Short loops mean more stable quadruplexes, while the relation between G-runs length and quadruplex stability is more complicated and cannot be described by a rule of thumb (Rachwal et al., 2007; Lightfoot et al., 2019(55,56)). Thus, when considering G4 motifs with four G-runs (which consist more than half of all bovine G4s), promoter and 3-UTR G4 motifs have longer G-runs and shorter loops than the average genome-wide (Fig. 3B). 5-UTRs and CDSs have longer than the average loops, while introns have longer than the average G-runs (Fig. 3B). We next focused in the patterns of G4 motifs based on their loop sizes. Starting with the G4s on promoters, in Fig. 3C it is clear that the G4 motifs with short loops are more frequent than those with longer. The above observation, which is already reported for other species (Lightfoot et al., 2019(56)), is almost the rule for all genic features for both template and coding strands (Supplementary Fig. 2). Exceptions are the coding strand of the 5-UTRs, in which pattern 6 is remarkably high and the template strand of the CDSs, for which the pattern is almost lost and the longer pattern (no 8) is the most frequent (Supplementary Fig. 2). The bias for shorter loops in all cases examined may reflect a need for thermodynamically stable quadruplex structures associated with genes. The differences in the pattern frequencies between the features could reflect functional differences linked to the roles that G4s play when located in each feature. The functional role of the G4s located on the template strand of CDS is as a negative regulatory effector on transcription elongation (Varshney et al., 2020(18)). The small number of rather non-stable G4s points towards an infrequent and transient regulatory effect on transcription.

### 3.3. SNPs on G4 motifs

The genomic regions that can adopt non-B DNA structures are known to be polymorphic (Du et al., 2014(57)). Regarding G4 motifs, this polymorphism can potentially affect their ability to form quadruplex structures (Zeraati et al., 2017(58)) and thus we next tried to explore the polymorphisms that arise from single nucleotide substitutions (SNPs) on the bovine G4 motifs. Consistent with the previous parts of the study, we focused on the SNPs that fall in the transcribed regions and their upstream regulatory areas. We obtained SNPs from three different databases. From the dbSNP database we obtained ∼99 million SNPs without any information about their frequencies, while from the Bovine Genome Variation Database (BGVD) we obtained ∼60 million SNPs along with their minor allele frequencies (MAFs) from 432 animals. Since many researchers use commercial arrays for studying SNPs, we also analyzed the ∼0.7 million SNPs from Illumina’s BovineHD BeadChip and we provide a list with all SNPs that fall on G4 motifs within transcribed regions and their upstream areas along with information regarding whether G4s are located on the coding or template strand (Supplementary Table 2).

SNPs can be on both G-runs and loops of G4 motifs. Among them the ones that fall on any of the following: a) any G in a 3-G-long G-run, b) the second or third G of 4-G-long G-runs and c) the middle G of 5-G-long G-runs, could disrupt the quadruplex structure (Nakken et al., 2009(38)). These SNPs hereafter are called disruptive SNPs (dsrSNPs).

By examining the difference in densities between dsrSNPs and non-dsrSNPs (SNPs that do fall on G4 motifs but do not disrupt the G4 structure) using the dbSNP, it is clear that disruptive nucleotides are less polymorphic (Fig. 4A; Supplementary Fig. 3). This is in line with what is reported for human G4 motifs (Nakken et al., 2009(38)). This lower degree of polymorphism, which implies a functional importance of quadruplex structures for all genic features, cannot be (at least not solely) attributed to mutability differences due to sequence context, since it is shown that at least in human 5’-UTRs and 3’-UTRs, the GGG sequences of G4 motifs are less mutated compared to the GGG sequences that do not overlap with G4s (Lee et al., 2020(59)).

**Figure 4:**
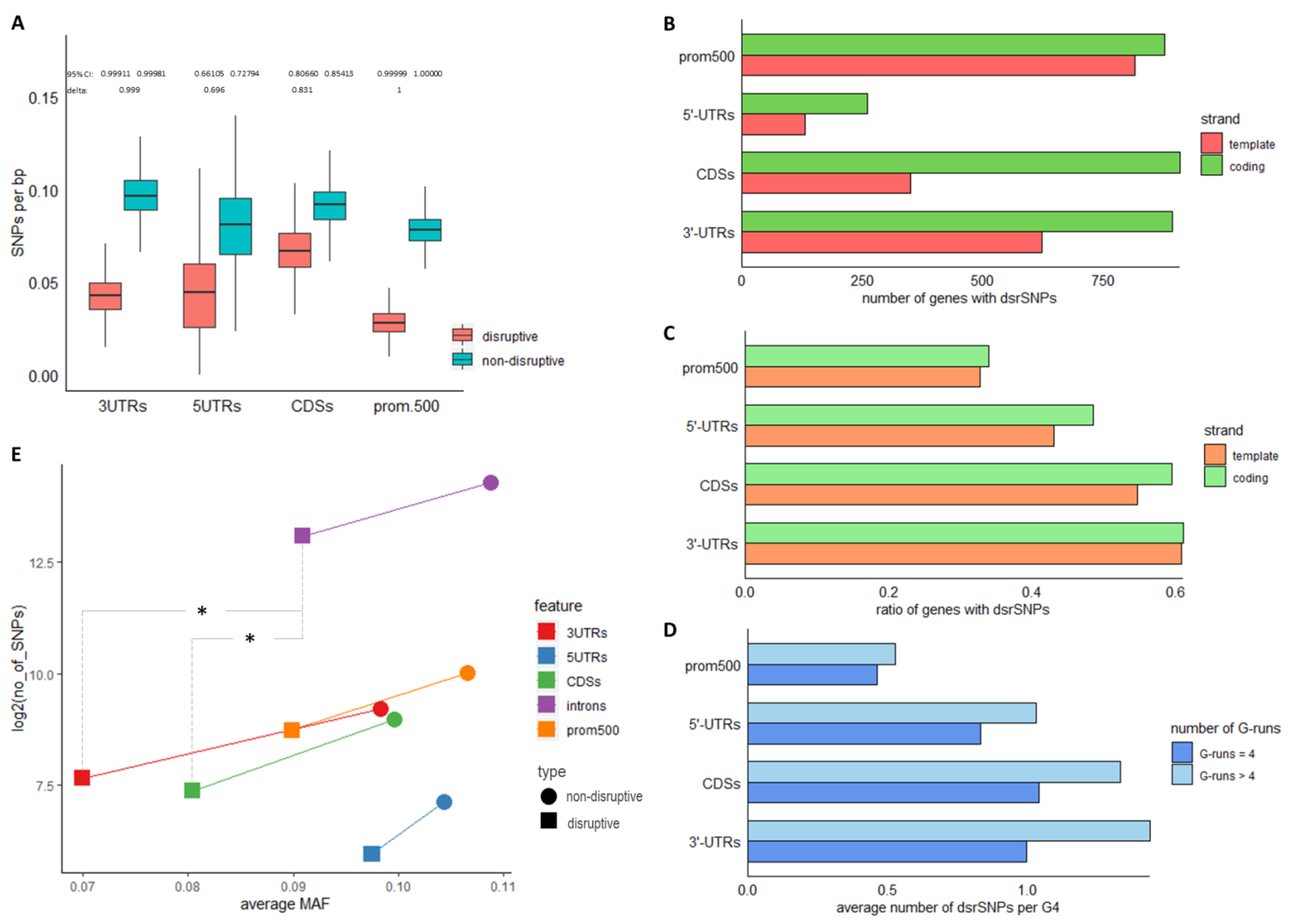
SNPs on bovine genic G4 motifs. **A**. Densities of disruptive and non-disruptive SNPs on genic G4 motifs of chromosome 28. 400,000 SNPs of chromosome 28 were randomly selected and the densities of the disruptive and non-disruptive SNPs on the G4 motifs were calculated. The randomization test was conducted 1,000 times. Boxes show the second and third quartiles. Whiskers extend up to the higher and lower values excluding the outliers, defined as values further than 1.5 fold the distance between the first and third quartile. Effect sizes were calculated using Cliff’s delta. SNPs source: dbSNP. **B**. Distribution of disruptive on genic features. The bars show number of genes with one or more disruptive SNPs on the G4 motifs that are annotated on their coding or template strands. **C**. Ratio of the genes with at least one disruptive SNP on the G4 motifs that are annotated on their coding or template strands. SNPs source: dbSNP. **D**. Average number of disruptive SNPs per G4 motif for motifs with 4 or more G-runs. SNPs source: dbSNP. **E**. Number and average MAFs of disruptive and non-disruptive SNPs on the G4 motifs of genic features. The Mann-Whitney U test was used to compare the MAFs among the features. *, p < 0.05. SNPs source: BGVD database with SNPs MAF ≥ 0.01.

By further assessing the dbSNP dsrSNPs on coding and transcribed strands we observed a high number of genes with dsrSNPs in promoters and a significantly lower one in 5’-UTRs (Fig. 4B). Furthermore, the differences between coding and template strands seem to mirror the ones of the number of genes with G4s (Fig. 1E). On the contrary, the ratio of genes with dsrSNPs (also dependent on the number of genes with G4s) has the lowest value for promoters and shows no obvious strand specificity (Fig. 4C). The existence of a high number of G4s on the promoters of many genes, with a considerable gene fraction to be free of dsrSNPs suggests that the integrity of the quadruplex structures must be important for this gene fraction.

When calculating the mutability of genic G4 motifs through the average number of dsrSNPs per G4, there is an apparent difference between the motifs with exactly four G-runs and the ones with more (Fig. 4D). Given the importance of G4s’ roles, the above result is expected since disruptive mutations in motifs with four G-runs would be deleterious for the quadruplex structure. On the contrary, the extra G-runs could act as “replacement parts” abolishing the catastrophic effect of a dsrSNP. When comparing the gene features, there is a correlation between the average number of dsrSNPs per G4 (Fig. 4D) and the average number of G-runs per feature (Fig. 3A). This means that a high number of G-runs per motif or a low average number of dsrSNPs work in the same direction regarding the tolerance against mutations. The length of G-runs is one more factor associated with mutability since the longer G-runs have less disruptive nucleotides. The consistency between the average number of dsrSNPs per G4 and length of G-runs is also present for promoters, 5’-UTRs and CDSs but it doesn’t seem to be the case for 3’-UTRs (Fig. 1E). Taken together, the above point towards a set of safeguards for the maintenance of the quadruplex structures’ integrity which is rather differential between the genic features.

Additionally to this “mutational stability” that is examined above, a kind of “structural stability” of the quadruplexes has been also described, where motifs with shorter loops tend to form more stable quadruplexes (Lightfoot et al., 2019(56)). For bovine genic G4 motifs it seems that the “mutational” and “structural” stabilities are correlated when taking into consideration the average loop lengths (Fig. 3B) and the average number of dsrSNPs per G4 (Fig. 4D). A correlation between loop-associated *in vitro* stability and conservation has been also reported for human G4s (Lombardi et al., 2019(60)).

As already mentioned, besides the SNPs from the dbSNP, we also analyzed those from BGVD. From the ∼60 million SNPs of the database, we took into consideration only the ones with MAF ≥ 0.01, and from these we considered only the ∼32K that are annotated on the genes and their promoters. Fig. 4E shows that in line with the dbSNP, the number of SNPs on disrupting nucleotides is lower than the number on the non-disruptive nucleotides for all genic features. The two databases are also similar when comparing the number of dsrSNPs among the genic features. The exception is the promoters, which have a high number of dsrSNPs in BGVD but low in dbSNP. This difference occurs also when BGVD SNPs are not filtered for MAF ≥ 0.01 (data not shown). Similarly to the number of dsrSNPs, the average MAFs of the disruptive positions are also lower than the non-disruptive. The reduction of SNPs/MAFs from non-disruptive to disruptive nucleotides is similar for all features except 5’-UTRS for which the SNPs number reduction is stronger compared to the MAF reduction. The disruptive nucleotides of CDSs and 3’-UTRs have the lowest MAFs among all features. This reduced MAF could be a signal of negative selection for the G4 motifs of these particular genic areas.

From the results shown in Fig. 4, it is apparent that the different genic features differ in terms of mutability and polymorphisms of their G4 motifs. This could be connected to the specific processes and functions that the G4s in each feature serve as well as the importance of these functions.

Given the functional involvement of G4s in several processes, the variability of genic G4s can be a valuable tool for better understanding their role as well as for genetic improvement by animal breeders (Stefos et al., 2021(30)). Fig. 5A shows that indeed there are QTLs/associations that colocalize with genic G4s. Although the overall distribution of the traits connected to the QTLs/associations colocalizing with G4s does not show any outstanding difference compared to the traits distribution of the whole database, all individual trait ratios, except for “reproduction”, differ significantly between the two QTL/associations sets (Fig. 5B). By the present study we do not aim to make any genetic correlations of G4 motifs with specific phenotypic traits, however we suggest that employing G4 motifs in genetic studies of cattle may serve as a valuable tool providing an extra layer of information.

**Fig. 5:**
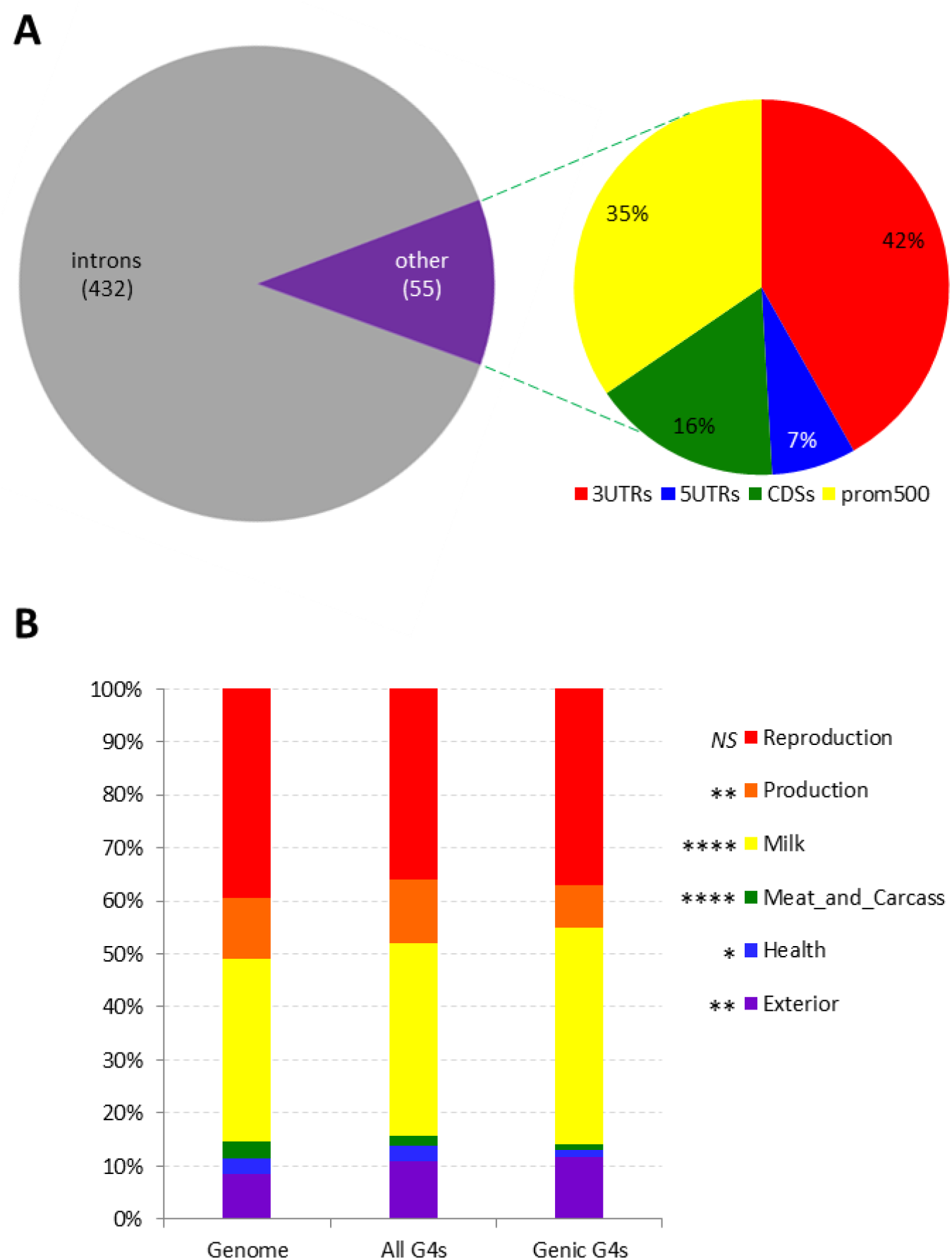
Bovine QTLs/associations of the QTLdb and G4 motifs. **A**. Proportion of genic G4 motifs that colocalize with QTL/associations. **B**. Proportions of the traits linked with the QTLs/associations: all QTLs/associations of the database (Genome), QTLs/associations that colocalize with genic and intragenic G4 motifs (All G4s), QTLs/associations that colocalize with genic G4 motifs (Genic G4s). The asterisks indicate significant differences for the representation of the traits between the genome QTL/associations and the genic-G4s. The significance was tested by randomly selecting from the genome QTL/associations a number of QTL/associations equal to the number of genic G4 QTL/associations, and then by comparing the traits ratios of the random sample to that in the genic-G4s QTL/associations. The randomization test was conducted 10,000 times. As p we define the ratio of the randomization tests with different (higher or lower) than the relative genic G4 value. NS, Non-Significant; *, p < 0.05; **p < 0.005; ***, p < 0.001 ****, p < 0.0001.

### 3.4. G4s and transposable elements

Trying to rationalize the occurrence of the loop and the G-run patterns on the genic features (Supplementary Fig. 2), we observed specific patterns to be overrepresented in introns. More detailed examination revealed that these overrepresented patterns are due to identical G4 motifs that are highly repeated within introns (Supplementary Fig. 4). Interestingly, the rest of the genic features do not carry repeated G4 motifs in as high relative frequencies as in introns.

Since a possible explanation for a given sequence to repeatedly occur in the genome is to be part of transposable elements (TE), we examined the association of G4 motifs and TEs. As it is shown in Fig. 6A, the colocalization of G4s and LINEs in introns is slightly higher than the expected and lower than that in the whole genome. For all the other genic features this colocalization is much lower than the expected. Oppositely to LINEs, for the other main orders of retrotransposons the likelihood for colocalization with G4s is considerably low (Supplementary Fig. 5). In Fig. 6B it is shown that in fact, among all genic features, only introns are considerably populated with LINEs. This is expected since the transposition of LINEs in other features, in CDSs for example, could be catastrophic. However, this “loose” consistency between LINE density and likelihood for co-occurrence with G4 motifs on genic features led us to further examine the LINE elements. An extra fact for going deeper into LINEs was that although their co-occurrence with G4 motifs has been already reported genomewide in other species (Kejnovsky et al., 2015(61)), to the best of our knowledge studies focusing in this co-occurrence within genes are missing.

**Figure 6:**
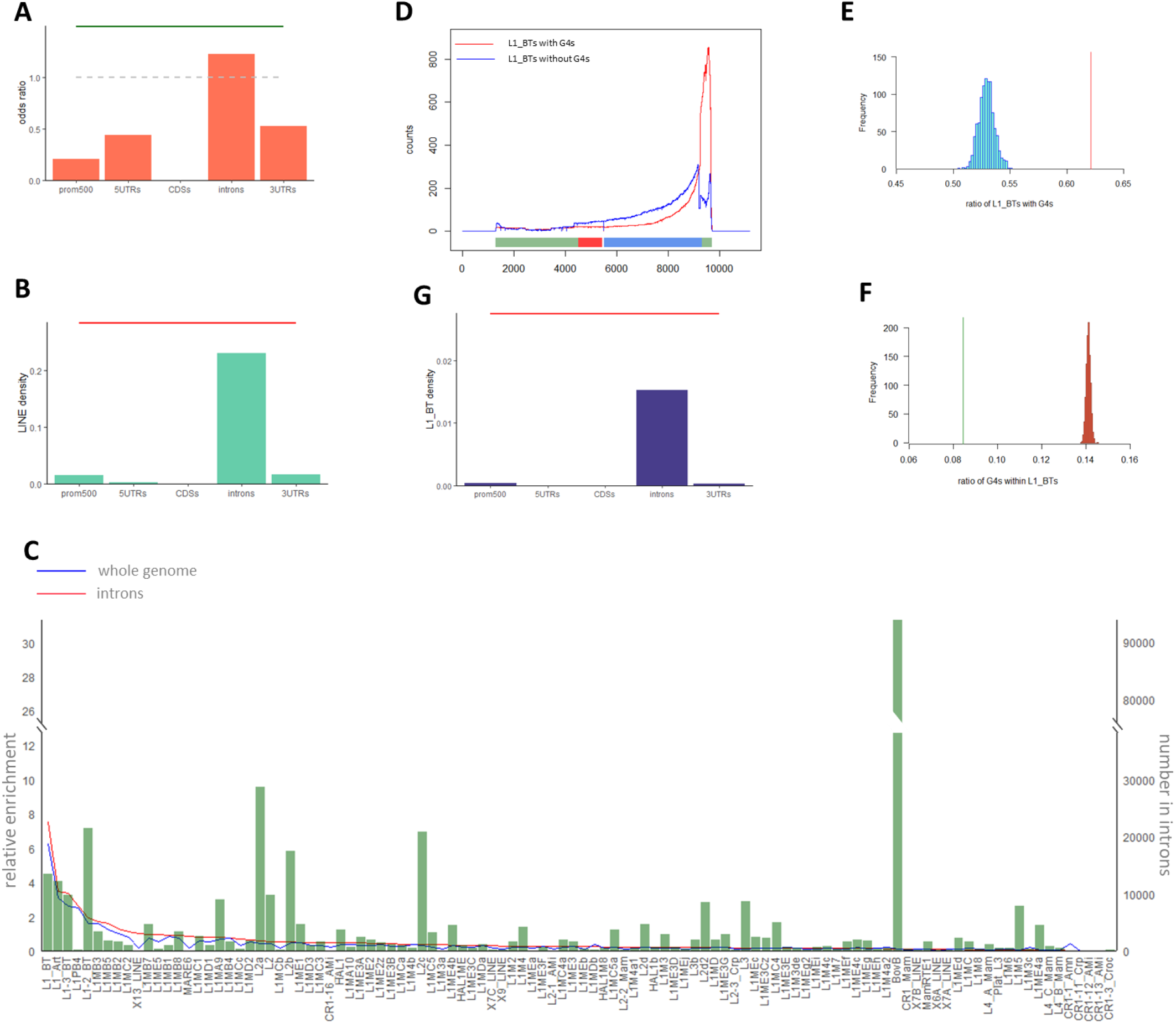
Overlap of LINE elements with bovine genic G4 motifs. **A**. Likelihood of G4s and LINEs colocalization in genic features expressed as odds ratios. The green horizontal line indicated the odds ratio for G4s/LINEs colocalization in whole genome. Odds ratios have been calculated as (length of areas with G4s-LINEs co-occurrence)*(length of areas free of G4s and LINEs)/(length of areas with G4s and free of LINEs)*(length of areas with LINEs and free of G4s). **B**. Density of LINEs in the genic feature. The density is calculated as the total length of the LINEs on a feature divided by the total length of the feature. The horizontal red line indicates the LINE density in the whole genome. **C**. LINE families with G4s. Bars show the absolute number of repeat elements of each family in introns (right y-axis). Lines show the relative enrichment of repeat elements in G4s in whole genome and introns (left y-axis). For calculating the relative enrichment, first the relative frequency of LINEs overlapping with G4 motifs was calculated and then was divided by relative frequency of each LINE family. For reasons of visualization, only the LINE families that overlap with G4 motifs in introns are shown on x-axis. **D**. Alignment of intronic L1_BTs with a full length element. 1000 random intronic L1_BTs that carry G4s (red line) or not (blue line) were aligned against a full length bovine L1_BT element that carries a G4 motif on its 3 UTR (GenBank: DQ000238.1). The lines indicate the number of times that each nucleotide of the full L1_BT is represented in the 1000 random L1_BTs. Above the x-axis, the features of the full L1_BT are shown: red for ORF1 (transposase); blue for ORF2 (endonuclease reverse transcriptase); grey-grey for non-coding sequences. **E**. Intronic and whole genome L1_BTs that carry G4 elements. The ratio of intronic L1_BTs that carry G4s is depicted by the vertical red line. The histogram shows the ratios of 1000 random L1_BTs lists sampled from the entire bovine genome and of size equal to the number of all intronic L1_BTs. **F**. Intronic and whole genome G4 elements located within L1_BTs. The ratio of intronic G4 elements that are located within L1_BTs is depicted by the green vertical line. The histogram shows the ratios of 1000 random G4s lists sampled from the entire bovine genome and of size equal to the number of all intronic G4s. **G**. Density of L1_BTs in the genic feature. The density is calculated as total length of the L1_BTs on a feature divided by the total length of the feature. The horizontal red line indicates the L1_BTs density in the whole genome.

Thus, the next step, given that LINEs include many families that may differ in how often occur within introns or overlap with G4 motifs, was to examine all these LINE families. As shown in Fig. 6C, L1_BTs are exceptionally enriched for G4s in introns compared to all the other families and L1-2_BTs and L1-3_BTs are included in the top five. L1_BTs are also enriched for G4 motifs also in the whole genome. The relative enrichments in introns and whole genome follow a similar distribution among the families without any remarkable exceptions (at least for the families with a considerable number of elements). L1_BTs are still active elements (Adelson et al., 2009(62)) and are among the youngest LINE-1 elements in cattle. This is in agreement with what has been described in humans, where G4 motifs are more enriched in younger LINE-1 families (Sahakyan et al., 2017(63)).

Since L1_BTs are significantly enriched in G4 motifs genomewide as well as in intronic areas, we considered the idea that these repeat elements act as vehicles for G4 motifs delivery in introns, in the same way that some human and plants LINE-1 families have been suggested to deliver G4s genomewide (Kejnovsky et al., 2015(61)). In these organisms LINE-1 elements are enriched for G4s in their 3’UTRs. Given that incomplete reverse transcription during the replication of LINE-1 often results in truncated elements at their 5’-end (Han 2010(64)), these elements could have the advantage to deliver short sequences carrying G4 motifs without the need to copy/paste the full length element (Kejnovsky et al., 2014(65)). In order to investigate the above model in cattle, we aligned intronic L1_BTs with a full length L1_BT element that carry a G4 motif on its 3’UTR. Indeed the vast majority of the intronic L1_BTs that carry G4s are truncated elements that are aligned on the G4-containing region (Fig. 6D). On the other hand, the intronic L1_BTs without G4s that were used as control, poorly overlap with the 3’-UTR carrying the G4 motif. It should be noted that the above observation is also the case for the non-intronic L1_BTs (data not shown).

We next focused in the comparison between the co-existence/overlapping of G4s and L1_BTs in introns and the whole genome. Regarding the ratio of L1_BTs that carry G4 motifs, we show that intronic L1_BTs carry significantly more often G4s than the L1_BTs in the whole genome (Fig. 6E). This could mean that selective mechanisms may have favored the transposition of L1_BTs that carry G4s against those without G4s in introns compared to the rest of the genome. On the other hand, the intronic G4s are less likely to be part of a L1_BT element than the average genomic G4 element (Fig. 6F). This could mean that, despite the seemingly significant role of L1_BTs in G4 delivery in introns, the enhanced G4 density of this region compared to the whole genome (Fig. 1B) must be rather attributed to other mechanisms not associated with L1_BTs delivery mechanisms.

Among the genic features studied here, only the introns are susceptible to L1_BTs transposition (Fig. 6G). Despite that the intronic transposition appears to be moderately reduced compared to that of whole genome, it seems that the L1_BT activity is more connected to G4s delivery in introns than in whole genome. The answer to the question what is the need for G4 delivery in the introns by L1_BTs may be connected with general roles that are attributed to G4s like the fine regulation of transcription, the speed of replication within introns and the presence of epigenetic marks (Lexa et al., 2014(66)).

## 4. Conclusions

The present study is the first attempt dealing with the description of the G4 motifs of the bovine genome, paying special attention to those motifs which are associated with genes. Livestock models have several particular advantages over rodents in translational research and thus their increased engagement as model organisms can benefit both these species and human (Roth and Tuggle 2015(23)). In this direction we tried to highlight the similarities/particularities of the bovine G4 motifs compared to the human and we tried to go further by suggesting a way that G4s are delivered in the bovine genes. In particular we showed that the distribution of G4s among the genic features is similar to what is already shown for human and the ontologies/pathways of the bovine genes carrying G4s show no surprises compared to other species. We describe in detail the structural characteristics of the bovine G4s and we point out that the link between structural and mutational stability is similar to that described for other species. We also show that similarly to humans, there are specific orders of bovine retrotransposons which are enriched in G4 motifs and we suggest one of them as a vehicle of G4 delivery to the introns of the bovine genes. To the best of our knowledge, although the role of repeat elements in delivering G4s has been already described, this is the first time that specific retrotransposons have been suggested for G4s delivery within genes. In parallel to the contribution in the field of G4s biology, our study can be also beneficial for the animal breeders and researchers in the farm animal field. We provide extended lists of bovine genes that carry G4 motifs for every genic feature as well as a list with all the G4-disrupting SNPs of one of the most used and well-established commercial arrays which in many occasions is routinely used in farm animals for genetic selection. We finally introduce the notion that there is a basis for linking traits of agricultural importance with the genetic variation associated with G4 motifs, underlining their potential in breeding. In conclusion, our work introduces the study of G4 quadruplexes in the research associated with livestock animals and serves as a starting point for deeper and more sophisticated G4s-associated works. The latter can serve towards rendering cattle as an interesting new model organism as well as in benefiting cattle research *per se*.

## Author contributions

Conceptualization was performed by GS, GT, IP; Manuscript was written by GS, GT, IP; Bioinformatics analyses were conducted by GS.

---

**Supplementary Fig. 1:**
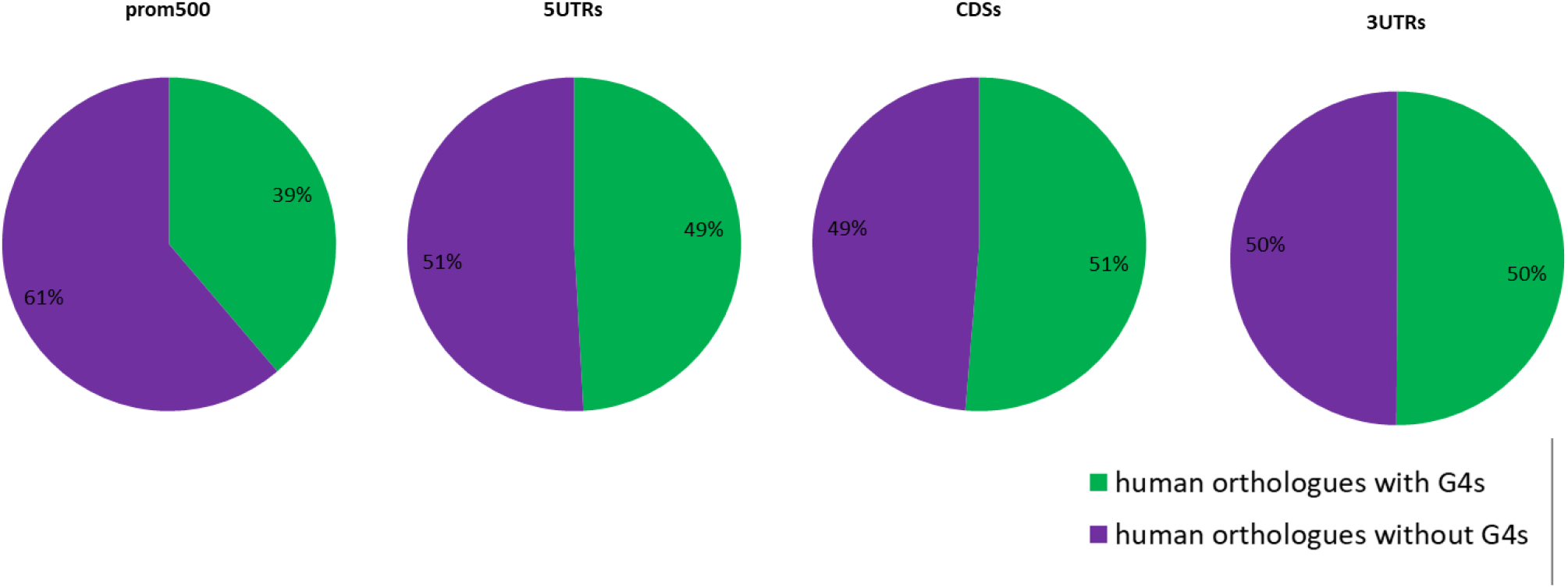
Occurrence of G4 motifs in human orthologues. The human orthologues of the bovine genes that carry G4 motifs are grouped to those that carry G4s in the same genic feature and those that do not.

**Supplementary Fig. 2:**
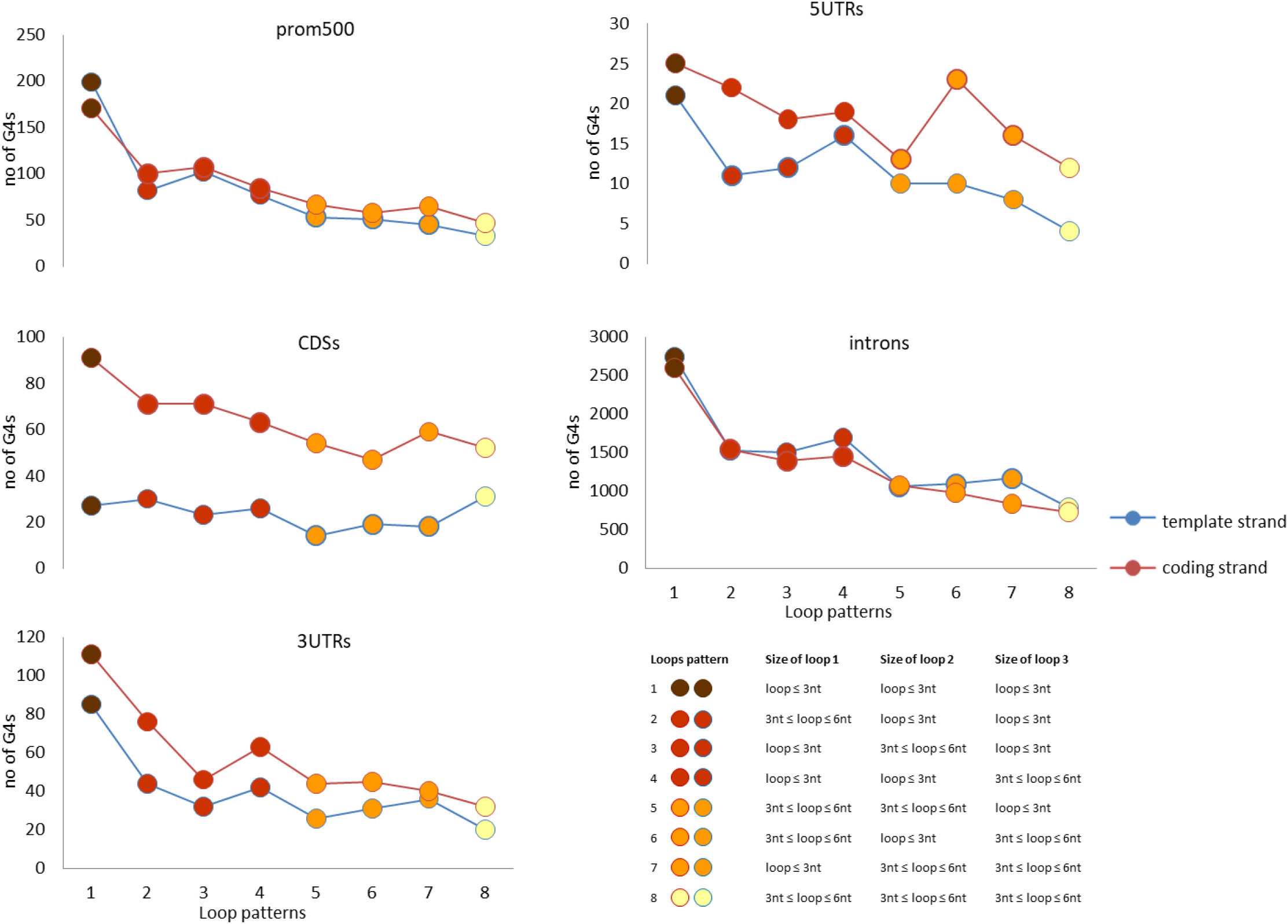
Number of G4 motifs annotated in 8 patterns according to the lengths of their loops. G4s with four G-runs and length of G-runs smaller or equal to 4 nucleotides were only taken into consideration. The description of the patterns is shown in the appendix at the lower right corner. The orientation of the motifs according to transcription direction was taken into consideration for the annotation to the patterns.

**Supplementary Figure 3:**
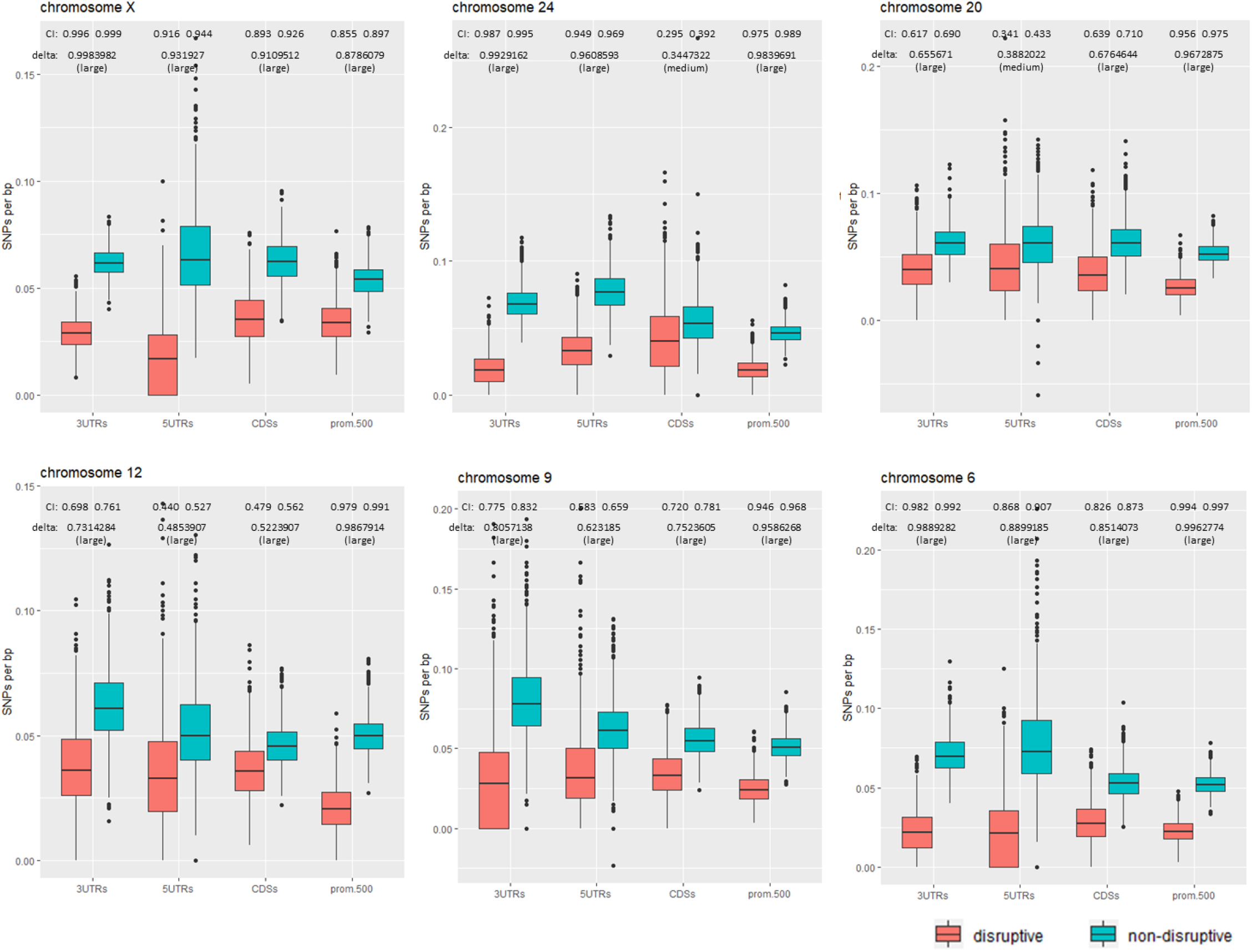
Densities of disruptive and non-disruptive SNPs on genic G4 motifs of several bovine chromosomes. For each chromosome, 400,000 SNPs were randomly selected and the density of the disruptive and non-disruptive SNPs on the G4 motifs were calculated. The randomization tests were conducted 1,000 times. Boxes show the second and third quartiles. Whiskers extend up to the higher and lower values excluding the outliers, defined as values further than 1.5 fold the distance between the first and third quartile. Effect sizes were calculated using Cliff’s delta. SNPs source: dbSNP.

**Supplementary Figure 4:**
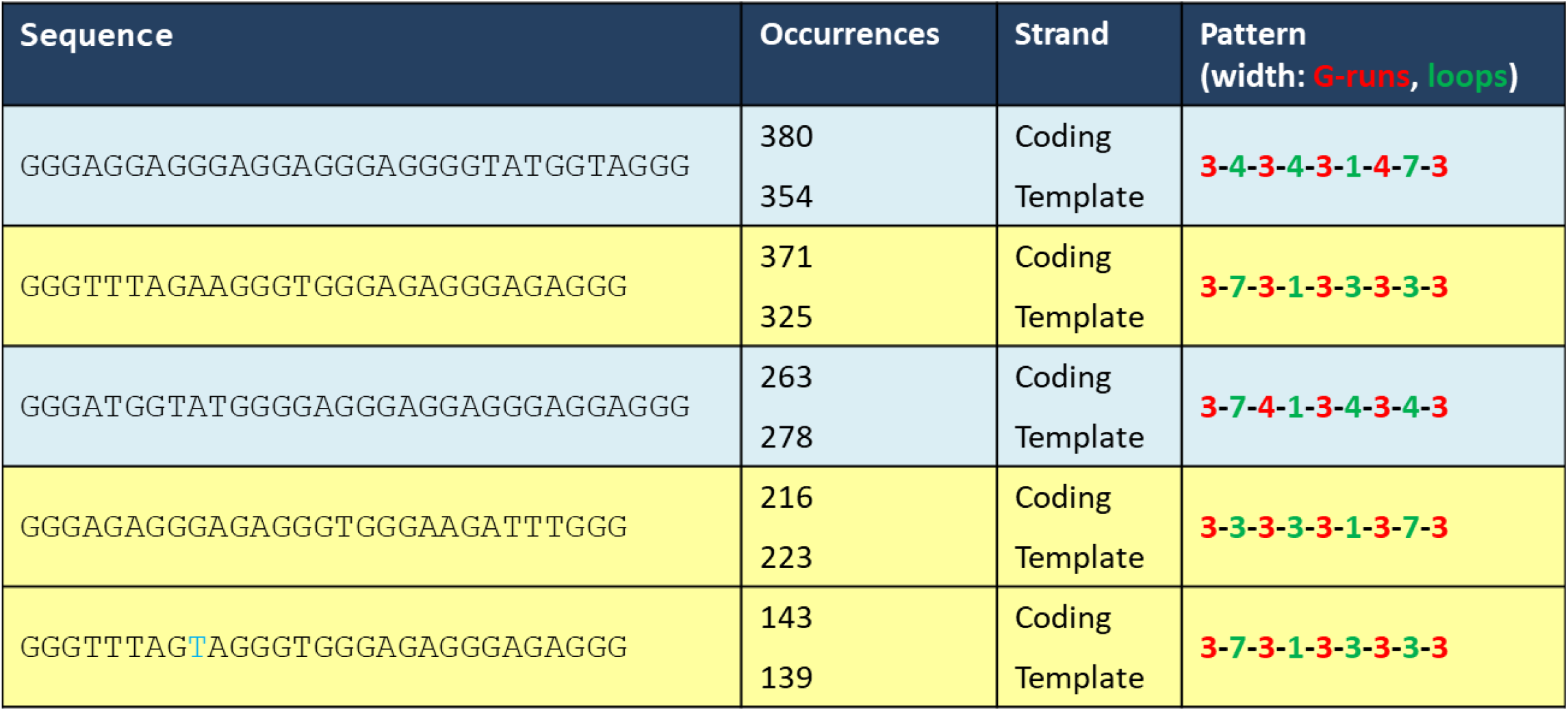
Top 5 of the more frequent G4 motifs found on introns. Motifs shown in the same background color are either the inverted versions of the same motif or variations differing in one nucleotide (indicated with blue).

**Supplementary Figure 5:**
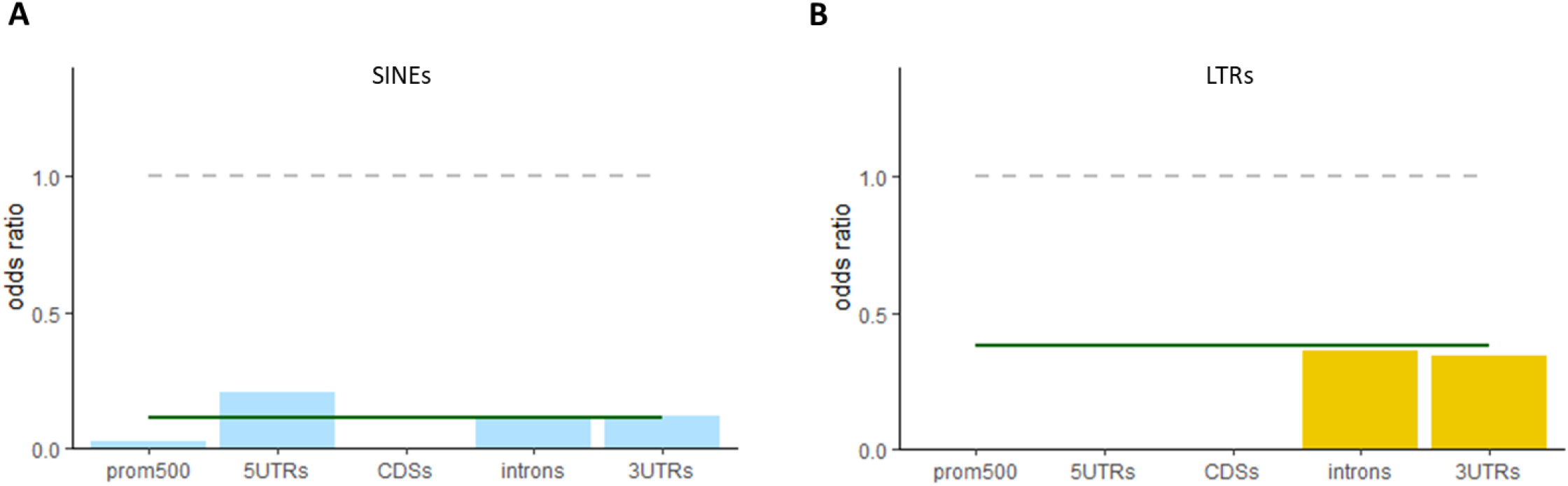
Colocalization of **(A)** SINEs and **(B)** LTRs with G4 motifs. The green horizontal lines indicated the odds ratios for G4s/TEs colocalization in whole genome. Odds ratios have been calculated as described in Fig. 6A.

